# Continuous ancestry profiles from genomes and genealogies

**DOI:** 10.64898/2026.11.11.765992

**Authors:** Gilles-Philippe Morin, Claudia Moreau, Simon L. Girard

## Abstract

Using a single population label to represent an individual’s ancestry may offer convenience, but ancestry is multidimensional, continuous, and reflects contributions from multiple ancestral sources. Here, we demonstrate empirically that ancestry profiles outperform categorical ancestry labels in capturing this complexity. Relatedness among 28,332 genotyped individuals was measured using haplotype painting and ordination and was regressed on regional ancestry proportions from the deep genealogies available for one-third of the cohort. When tested on participants in a validation set, predicted profiles placed 78% of documented ancestry in the correct region, against 48% for the training set average. Giving each participant a single regional label placed only 32% of their documented ancestry correctly, and a perfect classifier naming each person’s largest documented region would reach only 38%, because most individuals carry ancestry from several regions. Thus, the continuous profile captures substantially more ancestry than any classification, providing support for representing ancestry as a continuum rather than a label.

## Introduction

In population and medical genetics, it is commonplace to make use of an individual’s ancestry, whether to adjust for structure and stratification in association studies or to align participants with reference panels. This information is usually categorical and places a person in a cluster or assigns them to a given population. Such categories offer convenience, yet ancestry is not in itself categorical: it comes from genome segments inherited from many ancestors, who may or may not have a single origin.^1^ A single label reduces this admixture to its largest share, and increasing evidence suggests that discrete population descriptors misrepresent what is essentially continuous.^2,3^

Two types of methods estimate population structure without relying on labels, based on either allele-frequencies or haplotype. Allele-frequency approaches, such as principal components of a genotype relationship matrix (GRM) and model-based admixture estimates,^4,5^ are fast and scale to millions of samples, but they output axes that need interpretation. For instance, principal components can be a byproduct of sampling instead of reflecting a real correlation with geography,^6,7^ and additional tests are needed to isolate a real signal from noise.^8^ By contrast, haplotype-based methods identify segments inherited from common ancestors and detect structure at a finer scale than allele frequencies.^9,10^ This way, haplotype-based methods have distinguished populations at the level of a single county^11^ and a single founder community,^12^ and have improved correction for fine-scale structure in a national biobank.^13^

Neither type of methods have been validated on an external record of ancestral origin. In populations with genealogical registers, such a record is in fact available. In this case, a deep genealogy details how each ancestor is related to the individual and where they lived, which allows regional ancestry to be read directly instead of inferred. Founder populations with these registers, such as Quebec,^14^ Iceland^15^ and Finland,^16^ also show fine-scale genetic structure that follows the genealogy closely.^17^ Regional ancestry can thus be treated as a documented quantity to predict from the genome rather than a hidden quantity to infer. In livestock genetics, genomic ancestry proportions are compared with the ancestry composition derived from pedigrees.^18^ The same comparison has not been made for a human population at the scale of thousands of participants with deep genealogies. Although human populations having deep genealogies are rare, those without genealogies may still benefit from the validation and inference that the former can provide and rely on the population descriptors available.

We hypothesized that genome-wide relatedness retains each individual’s regional ancestry as a continuous profile, and this profile describes an individual’s ancestry more faithfully than a single regional label. Here we develop this ancestry mapping. Using haplotype relatedness among genotyped participants and regional ancestry proportions drawn from their genealogies, we fit a regression that estimates, for any individual, the expected proportion of ancestry from each region. Because the regression is learned once and then applied using genotypes only, it may be extended to thousands of participants who do not have genealogical data. Finally, we show that these regional ancestry profiles capture most documented regional ancestry, and that reducing the profile to a single regional label obscures most of an individual’s regional ancestry, because individuals are truly admixed.

## Material and methods

### Ethics

This research was approved by the ethics board of the Université du Québec à Chicoutimi (UQAC). All CARTaGENE participants provided written informed consent.

### Data sources

The CARTaGENE cohort,^19^ launched in 2007, comprises 43,032 individuals (aged 40 to 69 at initial assessment) who were recruited in Quebec’s largest cities.

Genotype data of 29,337 participants was obtained from one of four arrays: Omni2.5, GSAv1, GSAv2, and GSAv3.^19^ Each dataset was processed separately using PLINK v1.9,^20^ keeping participants with at least 95% SNP coverage. At the variant level, we kept autosomal SNPs with a call rate of at least 95% and in Hardy–Weinberg equilibrium (*p* > 10^−6^, one test per dataset). The four datasets were then merged into a final dataset of 140,402 autosomal SNPs for 28,332 participants.

Genealogical records were obtained from the Balsac population register,^21^ which connects Catholic parish records for Quebec’s French-Canadian population beginning in 1621. Among genotyped CARTaGENE participants, 9,405 have a Balsac identifier, with family lineages reaching up to 15 generations. The regions align with Quebec’s administrative divisions used in earlier work.^22^

### Overview of the inference pipeline

The steps below turn a participant’s genome into a predicted ancestry profile, from which a single region of origin is derived for comparison purposes: a summary of the proportion of the genome participants have in common, an ordination of that summary, a genealogical reference to fit against, and a regression joining both (Figure 1).

**Figure 1.**
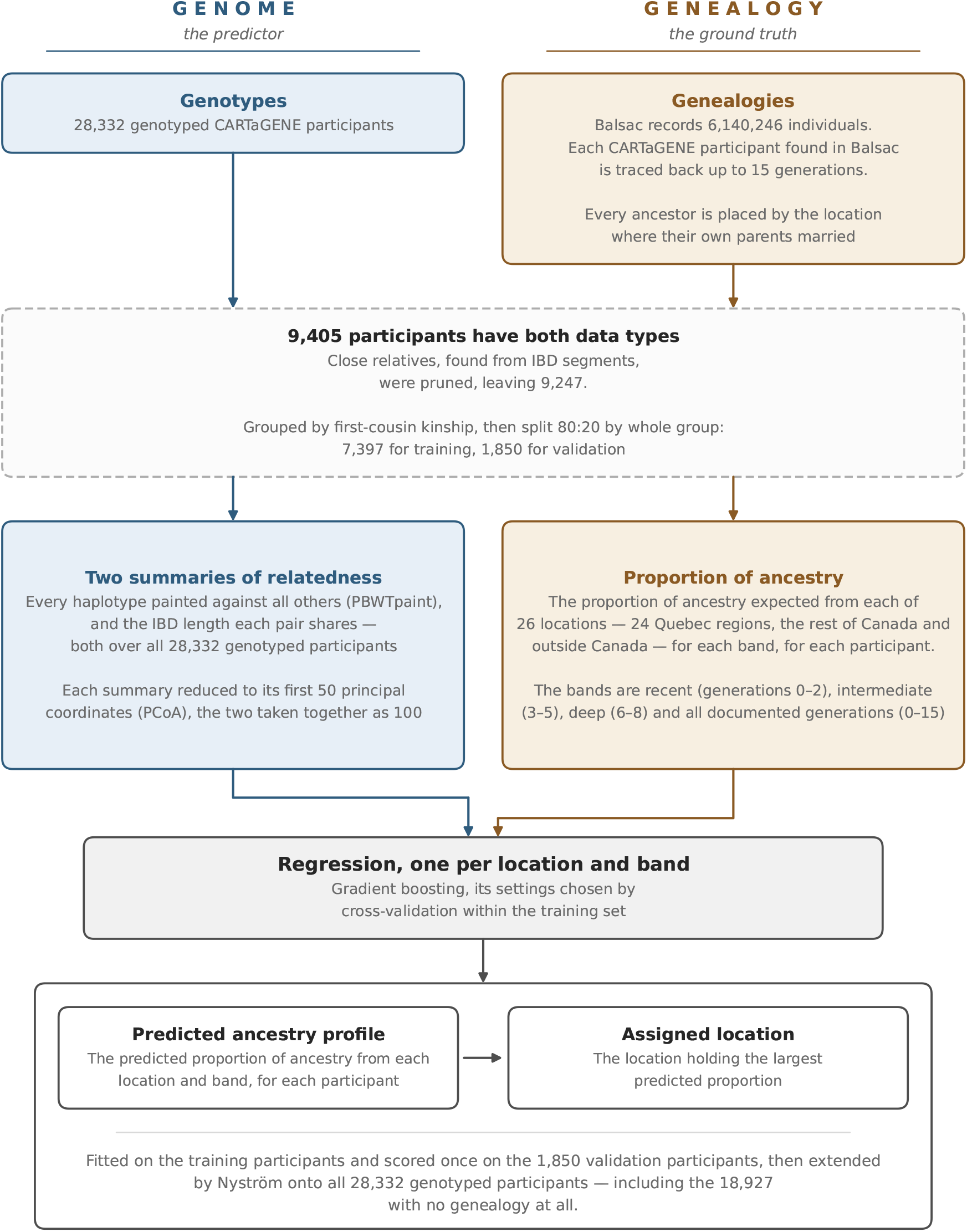
The regional ancestry inference pipeline.

### Two summaries of genomic relatedness

Before fitting any model, all genotypes must be condensed into a summary of participant relationships. A combination of haplotype painting and logarithm of observed kinship was chosen among eight summaries of the same genomes (see supplemental material and Figure S1).

Briefly, the coancestry matrix derived from the number of chunks during haplotype painting depicts each haplotype as a mosaic of the others under an approximation to Li and Stephens’ copying model.^23^ Painting was performed with PBWTpaint,^24^ implemented in pbwt on the positional Burrows–Wheeler transform,^25^ which paints every haplotype against all others in near-linear time. All 28,332 genotyped participants were painted chromosome by chromosome, the resulting chunk counts were summed over the 22 autosomes, and the matrix was made symmetric by averaging it with its transpose.

As for the observed kinship, genotypes were phased using Beagle version 18May20.^26^ Identity-by-descent (IBD) segments were identified with Refined IBD version 17Jan20^27^ at a LOD threshold of 3.0 and a minimum length of 2 cM, then merged using the merge-ibd-segments 17Jan20.102 tool. Observed kinship was derived from the total IBD length per pair across retained segments using the ibd_relatedness program by S.R. Browning (see *Web resources* below), as previously reported.^17^

### The training–validation split

To prevent bias from family structure, two relatedness thresholds were applied to the raw matrix of observed kinship. First, participants with *φ* ≥ 0.0884 (which corresponds, for example, to an uncle-niece relationship^28^) were removed. Removing such close relatives prevents the ordination from shifting principal coordinates toward family structure instead of population structure.^8^ This removal left 9,247 participants with both genotype and genealogical data. Afterwards, the remaining individuals were grouped using a threshold of *φ* ≥ 0.0442, which corresponds, for example, to a first cousins relationship. The resulting groupings were split into 8,766 families, most of them consisting of a single participant, with the largest containing five. Every group was assigned to the same side of the split to prevent bias from close relatives being found in both the training and validation sets.

Training and validation sets were split using a 80:20 ratio, with 7,397 participants in the training set and 1,850 in the validation set. All tests and metrics reported below were performed on the validation set. Entire families were assigned to one group or the other, and the split was stratified by region (see *Regional ancestry proportions defined from genealogies* below). Each region was represented in both sets in proportion to its size in the cohort. The validation set was excluded from fitting, tuning hyper-parameters, setting thresholds, and calibration. It was used to benchmark once, after all decisions were made.

### Ordination of the summaries of genomic relatedness

Summaries such as haplotype painting and logarithm of observed kinship provide one column per participant, which is far too many dimensions for a regression. Ordination by principal coordinate analysis projects those columns into a smaller space that preserves the distances between participants. The principal coordinates are then used to predict the regional ancestry profiles. For haplo-type painting, distances among training participants were derived from the symmetrized coancestry matrix of the number of chunks (*S*) using *D_ij_*^2^ = 2*c* − 2*S_ij_*, where *c* is the largest entry of *S*. The same formula was used for the logarithm of observed kinship. Classical multidimensional scaling computed the coordinates through eigen decomposition of the doubly centered matrix of squared distances.^29,30^ The validation set and genotyped participants without genealogy were projected in the same space via Nyström extension.^31^ This step adds a point to an existing principal coordinate diagram.^32^ One hundred axes were kept: 50 from haplotype painting and 50 from observed kinship.

### Regional ancestry proportions defined from genealogies

Population structure is usually described using the participants’ recruitment location or birthplace as an indirect indication of geographic origin. In our case, the genealogies provide these origins directly with an additional time component. The genealogies were processed with GeneaKit v0.1.0.^17^ For each person, the genealogy begins at generation 0 (the participant) and extends back to a maximum of 15 generations. Each occurrence of an ancestor at generation *g* has an expected genetic contribution of 2^−*g*^ to the participant’s genome.^33^ This ancestor is assigned to the region where their parents married and used as an indication for origin. Ancestors who do not have a documented marriage location in the register remain unassigned to a region. Every path to an ancestor is counted separately, so if an ancestor can be reached by two lineages, they are counted twice, and an ancestor and their parents each count at their respective generation. This procedure produces an ancestry profile for any region and generation. For a given region, a participant’s regional proportion is the sum of the expected genetic contribution of their ancestors from that region, divided by the total expected contribution of all documented ancestors. This total expected contribution corresponds to genealogical completeness, which at each generation equals the percentage of possible ancestors (2*^g^*) that are present in the population register.

Out of the 25 Quebec regions in the Balsac population register, a region was kept for training if at least 50 participants had at least 5% of their documented ancestry in this region in at least one of the bands defined below. These conditions were tested on the whole analysis set with both genomic and genealogical data before the training–validation split, to ensure that the same regions were modeled on both sides. One region (Nord du Québec) fell below the threshold and was excluded. Ancestry recorded outside Quebec was kept as two separate locations: the rest of Canada and outside Canada. Together these make 26 locations: 24 Quebec regions that pass the thresholds, the rest of Canada, and outside Canada.

Regional ancestry proportions were gathered using two approaches. The first approach collects all documented generations, 0 to 15, into a single measure that ignores the time component. The other approach splits the genealogies into three time bands: recent (generations 0–2), intermediate (3–5) and deep (6–8). These bands were chosen to see how ancestry can evolve over time through migration and admixture. Indeed, a region may be strongly represented in either recent or deep generations but not both, and a single measure would flatten these patterns. Using bands instead of single generations reduces the offset caused by uneven pedigree depth, and three bands of three generations proved adequate. No band covers generations 9 to 15, which are taken into account only for the first approach, because completeness drops sharply beyond the eighth generation and the record becomes mostly European. A deeper band would thus capture register limits more than provincial structure. Quebec regions contribute unequally to a participant’s ancestry (Figure S2) and enter the provincial structure at different generations (Figure S3). We can therefore trace the contribution of different regions at different time bands.

Since a band can be predicted only if the underlying record is robust enough, two criteria were applied to each participant and band. First, a participant’s band remains undefined if its ancestry completeness drops below 0.15, preventing reliance on a small set of documented lineages. Next, for a given participant, a band is also undefined if fewer than 25% of their documented ancestors in the band have a recorded region, since that band would otherwise be overshadowed by ancestry with no recorded place. Both thresholds were fixed before the split and used consistently for training and validation sets. Only participants who passed both tests received a score in that band. Among the 1,850 validation participants, 1,830 are scored on the recent band, 1,637 on the intermediate, 1,572 on the deep and 1,654 on the pooled band.

### Regression of ancestry proportions on genetic structure

The ordinations of haplotype painting and observed kinship are used together to predict the regional ancestry proportions read from the genealogies. One regression model was fitted for each region– band combination. There are four bands: the three generation bands defined above, and a flat band comprising all documented generations from 0 to 15. Every regression used histogram-based gradient boosting,^34,35^ implemented in scikit-learn 1.8.0.^36^ This non-linear technique can work with a hundred correlated coordinates without transformation.

Cross-validation was used during training: the training set was divided into equal parts, and at each training iteration a single part was left out to be predicted from the others. This way, every training participant had a prediction from a model that was not trained on them. Relatives always stayed in the same part. This partition was done twice: 1) over three parts, the settings of gradient boosting were tuned for each location and band, chosen from sixteen combinations of learning rate, number of trees, tree size and regularization. 2) over five parts, the partition produced new predictions using the previous tunings. Those predictions were scored by the coefficient of determination (*R*^2^), computed by location and averaged so that each location counts equally. This average is what we call the macro-*R*^2^.

The fitted regressions were then applied to all 28,332 genotyped participants using the same projection. Each regression returns a predicted profile for all genotyped individuals, which is the proportion of ancestry it assigns to each location in each band (Figure S4).

### Assigning a single predicted region to every participant

If classification is needed, the regional ancestry profiles have to be narrowed to one single location. However, systematically choosing the region with the largest proportion would tend towards the locations that are common in the training set, because a common location is more likely to be associated with a higher proportion for everyone. Each predicted proportion is therefore divided by that location’s mean proportion in the training set, raised to a power between 0 and 1. At 0 the largest proportion is systematically chosen, and at 1 the locations are ranked by how much they exceed their own average. The power was set from the training predictions, among 0, 0.25, 0.50, 0.75 and 1, as the one doing best, at the same time, on plain accuracy and on accuracy averaged over locations, so the rule cannot succeed on common locations only. The power was 0.50 for all bands. The rule assigns a location to every genotyped participant and counts as correct when this location holds the largest documented proportion in the band being scored.

### Validation and statistical analysis

Each reported metric, accuracies, *R*^2^ values, and *F*_1_ scores (the harmonic mean of precision and recall for one region in one band), comes from the validation set. None of the participants has relatives in the training set. Confidence intervals used bootstrap resampling of relative groups (1,000 draws),^37^ while p-values relied on permuting the participants’ documented regional ancestry proportions across equal-sized relative groups (10,000 permutations).

### Visualization

To assess whether the genetic structure appears without labels, the 100 principal coordinates of the haplotype painting and logarithm of observed kinship were mapped to two dimensions using UMAP 0.5.12, the Python version described by McInnes et al.^38,39^ The layout was initialized from the first two principal coordinates of that ordination, applying a minimum distance of 0.5 and 15 neighbors.

## Results

To predict regional ancestry from the genome, the haplotypes of 28,332 genotyped CARTaGENE participants were painted against each other, then this painting and the individual’s observed kinship logarithm were each reduced to 50 principal coordinates. As a reference, regional ancestry proportions were retrieved from the genealogies of 9,247 participants. To predict these ancestry proportions from the principal coordinates, gradient boosting regression was trained from a subset of 7,397 participants before extending the predictions to all 28,332 participants. Finally, the predictions were validated on the 1,850 remaining participants with both genotype and genealogical data who were excluded from model training and are unrelated to the training-set individuals (Figure 1). The results from this validation are reported below.

### Regional ancestry profiles are recoverable from genome-wide haplotype relatedness

The regression estimates ancestry proportions for every region, so each participant receives a distributed ancestry profile rather than a single label (Figure S4). A single regional assignment is derived afterwards from the region holding the largest predicted contribution.

Across all documented generations, the profile predicted for a validation participant overlaps the profile documented by their genealogy by 78% on average (95% CI 77.5–78.6). Overlap is the proportion of the documented profile matched by the prediction. For example, for a participant documented as 80% Charlevoix and 20% Mauricie but predicted as 20% Charlevoix and 80% Mauricie, the prediction matches 20% in Charlevoix and 20% in Mauricie, so the overlap is 40%. An ancestry-blind baseline with no genetic information, which gives every validation participant the mean genealogical profile of the training set, overlaps by only 48%. Adding genetic information halves the proportion of documented ancestry placed in the wrong region. Documented and predicted proportions correlate (*R*^2^ = 0.78) across the full generation range and across every region (Figure 2).

**Figure 2.**
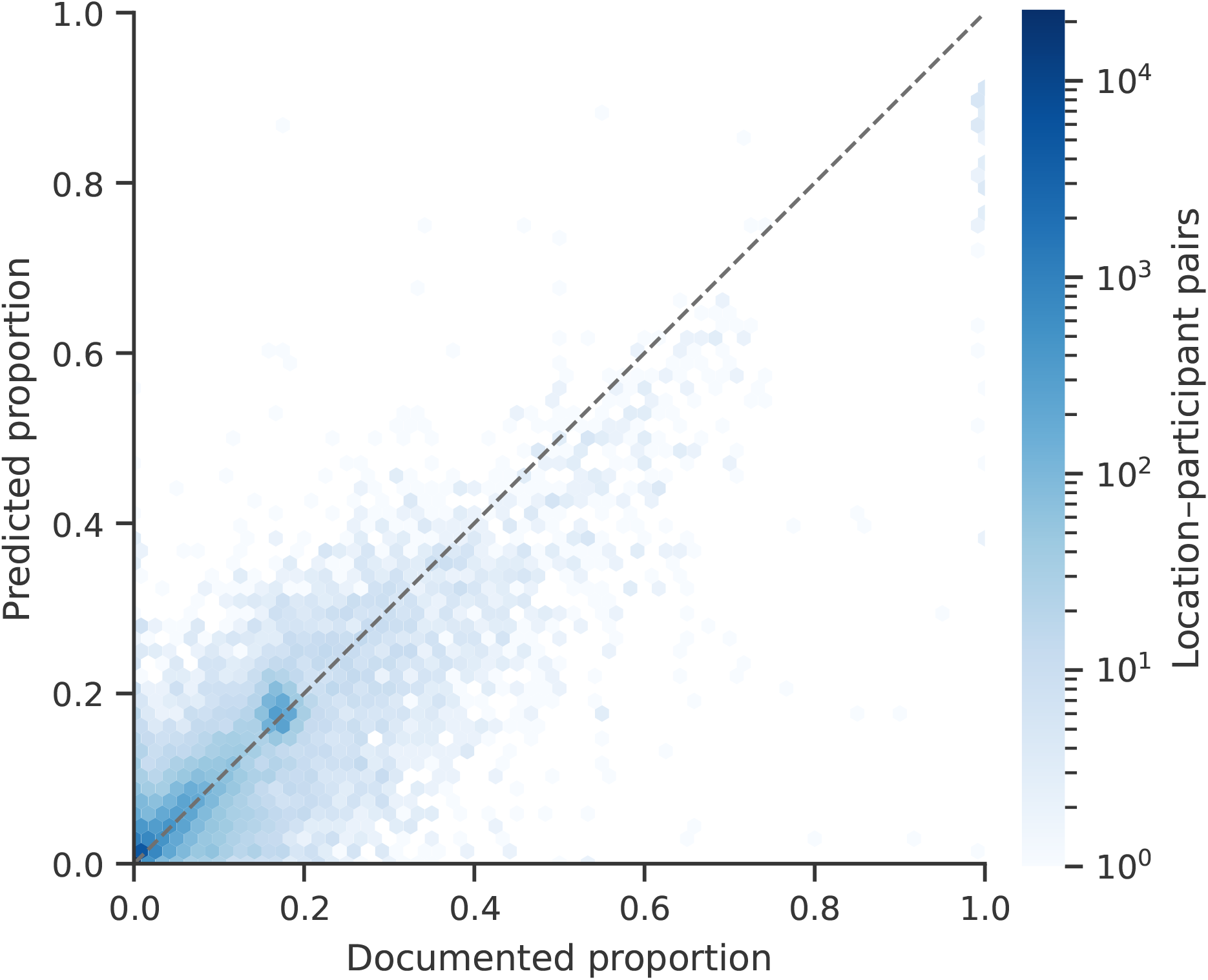
Predicted ancestry profiles reflect the documented ones. Each point represents the documented (horizontal axis) and predicted (vertical axis) ancestry proportions for one validation participant in one of 26 locations (1,654 participants × 26 locations = 43,004 comparisons). Because many points overlap, values are displayed as hexagonal bins, colored according to the number of observations on a logarithmic scale. The dashed line indicates equality between documented and predicted ancestry proportions. The overall coefficient of determination was *R*^2^ = 0.78.

### Accuracy of the assigned single label

To check whether a single region would be enough to accurately represent someone’s ancestry, the location with the largest predicted proportion was extracted for each genotyped participant after adjusting for how common that location is in the training set (Material and methods).

Classification using only the leading predicted region obscures underlying regional admixture. A single assigned region placed only 32% of a participant’s documented ancestry in the right place (95% CI 30.6–32.7). Even a perfect classifier, that would name each participant’s single largest documented region every time, would correctly place only 38% (95% CI 37.3–39.0). This is less than the 48% of the ancestry-blind profile. A label reduces a person’s ancestry to a single region, so only the proportion documented in that region counts toward accuracy. Conversely, the ancestry-blind profile spreads the average ancestry over all regions and accumulates partial accuracy in every region where the person does have documented ancestry.

The accuracy of the single label varies across the province (Figure 3). Overall, the assigned region is correct for 60% of validation participants. This participant-level accuracy, defined as the proportion of participants whose top predicted region matches their top documented region, differs from the proportion of documented ancestry correctly assigned by a single regional label. This means that 60% of individuals are assigned to the correct region, even though that region accounts for less than 40% of their ancestry profile. A baseline classifier that always predicts the most common region, Île-de-Montréal, would achieve only 20% accuracy. Of note, ancestry from areas isolated geographically or historically, such as Îles-de-la-Madeleine and Gaspésie, is well recovered, whereas neighboring regions of the Montreal urban area are often confused with one another. Smaller or more recently settled regions, such as Abitibi and Témiscamingue, were the least accurately recovered.

**Figure 3.**
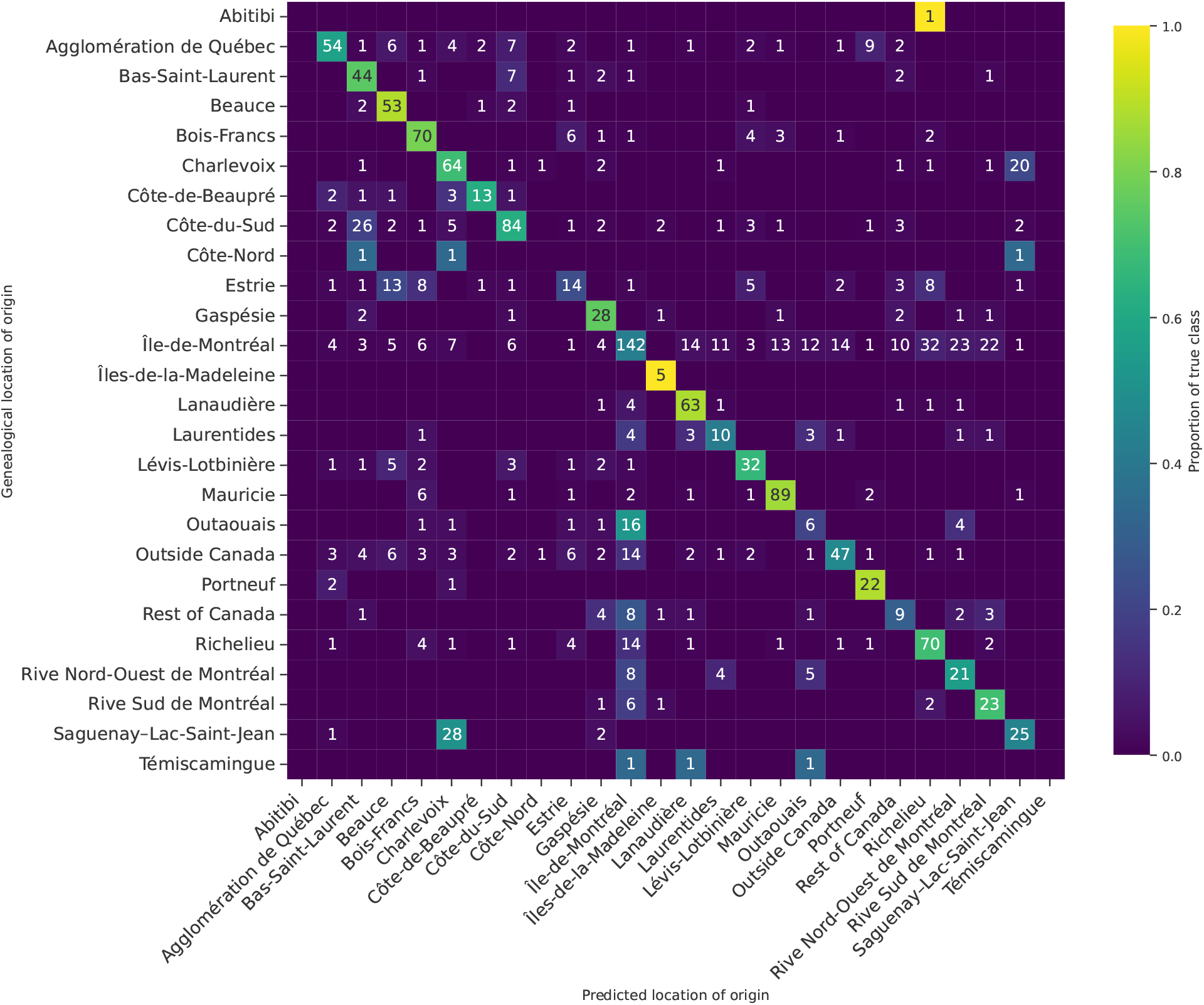
Accuracy of the predicted region of origin. Row-normalized accuracy of the region of origin predicted from genomic relatedness, against the documented region, across generations 0 to 15, among the validation participants.

To observe the evolution of regional ancestry profiles over a timescale, expected genetic contributions were split into three genealogical bands (0–2, 3–5, 6–8) in addition to being computed on the whole genealogy (generations 0–15, see Material and methods). Recovery depends on genealogical depth (Table S1). The deep band places the largest proportion of documented ancestry in the correct region, 83% against 55% for this band’s ancestry-blind baseline (95% CI 82.3–83.0, permutation *p* < 0.001. This band also has fewer documented origins. Only 16 locations are the largest documented region of at least one validation participant in the deep band, compared with all 26 in the recent band (Figure S5). Because the ancestry-blind baseline varies across bands, each band is best evaluated by the gain above the baseline toward a perfect profile rather than by the raw score itself. The intermediate and deep bands recovered regional ancestry proportions with similar accuracy, both outperforming the recent band. The band yielding the best region recovery varies by location (Figure S6).

### The UMAP projection

The structure captured by these models exists before any model fitting. Participants cluster when projected into two dimensions, even though the projection is built from haplotype relatedness alone, without any regional information as input (Figure 4). The single regions predicted by the genome and the regions documented by the genealogy largely coincide. A group of participants who are documented as coming from outside Canada splits from the main French-Canadian cluster and is largely of unrecorded origin. Regions also overlap within individuals: coloring the projection by the predicted proportion of one region at a time shows that most participants carry ancestry from several regions (Figure S7).

**Figure 4.**
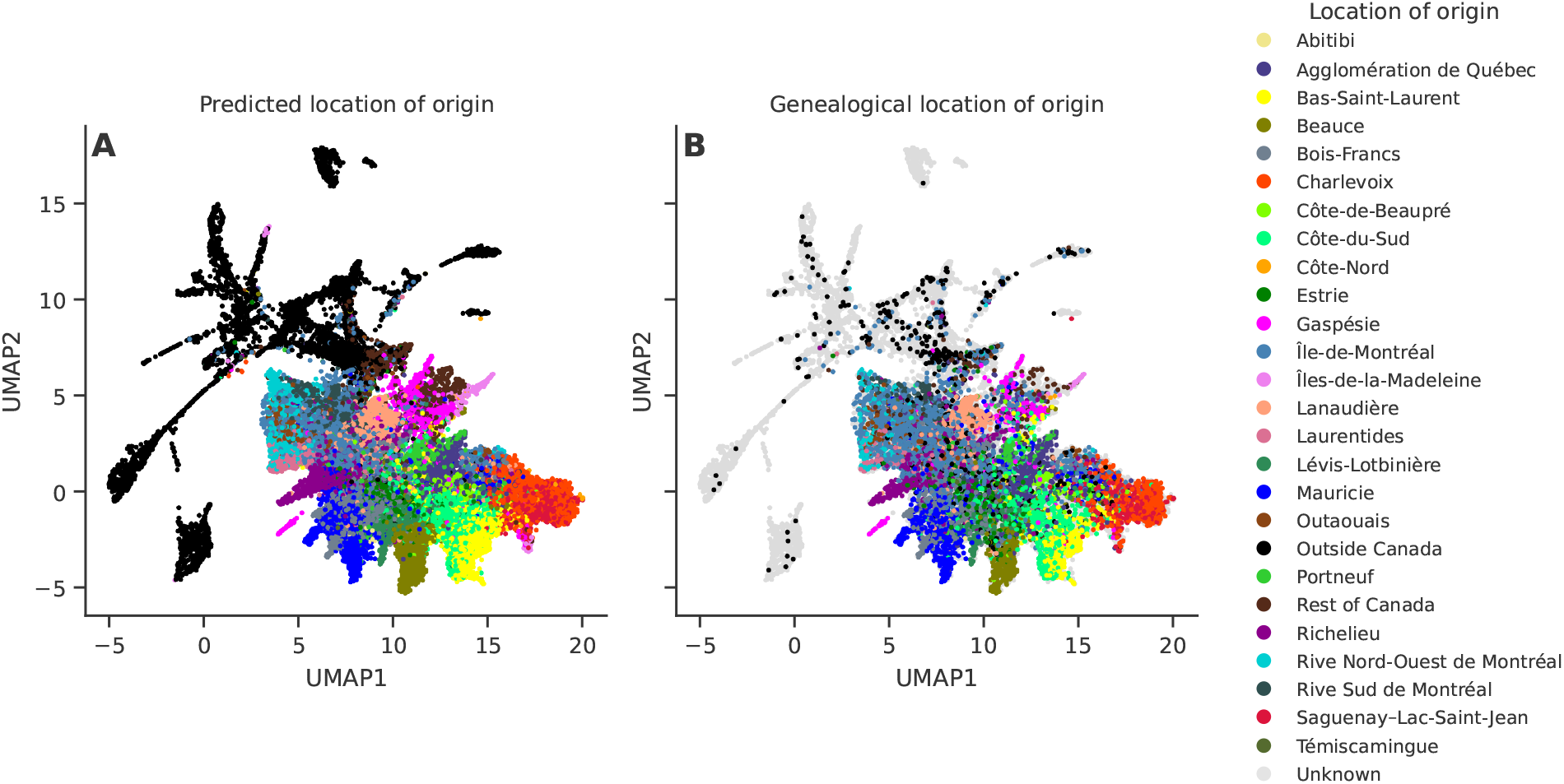
The genetic projection recovers genealogical structure. Both panels show the same UMAP of 28,332 CARTaGENE participants, built from the 100 principal coordinates of the combined summary of genomic relatedness. Only the coloring differs. Each person is colored by their predicted (A) or documented (B) region of origin (documented for the 9,405 individuals with genealogies), the region being the one holding the largest proportion of their placed ancestry.

### The recruitment site is a coarse approximation

A final check uses no genealogical information. CARTaGENE recruited in a small number of urban sites, dominated by Montreal, so the recruitment site provides only a coarse approximation of regional ancestry. In fact, Montreal is the most common recruitment site both for regions where participants’ parents married (Table S2) and for predicted regions (Figure S8). Nevertheless, predicted regions with their own recruitment site, such as Saguenay–Lac-Saint-Jean and Mauricie, are enriched among the participants recruited at the matching site. Because recruitment information is not used by the model, this partial correspondence provides independent validation, and the limited resolution of recruitment as a label further supports the use of genealogical profiles.

## Discussion

As genetic studies increasingly include individuals with mixed and diverse ancestries, recent work has highlighted the limitations of discrete ancestry labels and the value of modeling ancestry as a continuum.^2,3,40^ This perspective is reflected in recent work showing that prediction accuracy, polygenic scores, and association analyses vary along an ancestry continuum rather than between discrete ancestry groups.^41–43^ Here we show that genome-wide relatedness captures an individual’s regional ancestry as a continuous, quantitative profile rather than a category. Indeed, the predicted profile places 78% of a participant’s documented ancestry in the correct regions, far above a cohort-average baseline. Although a single assigned region is a convenient summary, it is not a faithful description. Even a perfect label, that identifies each participant’s largest documented region every time, accounts for only 38% of their documented ancestry which falls below the 48% achieved by a cohort-average profile, highlighting that a single region captures only about a third of a person’s ancestry on average. This suggests that the limitation lies primarily in the multidimensional nature of ancestry rather than in the performance of the classifier itself.

This observation raises a broader question of how ancestry should be represented and measured. Model-based admixture methods summarize ancestry as proportions of latent components,^4,5^ whereas local ancestry methods assign genomic segments to reference populations.^44–47^ Deep genealogical records provide a different source of information, based on documented ancestral contributions rather than genetic inference alone. In our study, the availability of such genealogies offered the opportunity to evaluate continuous ancestry against an independent historical record. More generally, such records can provide a valuable tool for assessing and interpreting ancestry profiles.

Our access to genealogical records also made it possible to examine how ancestry changes across generations. Migration events can be captured by the different generational bands of ancestry profiles but are lost when ancestry is summarized as a single estimate. The Charlevoix-Saguenay–Lac-Saint-Jean (SLSJ) recent migration provides an example of this phenomenon, with recent ancestors concentrated in SLSJ and more distant ancestors concentrated in Charlevoix. A single label of ancestry would therefore mask the movement of ancestors across regions over time. Examining ancestry by generational bands may therefore offer a useful approach for identifying migrations more generally, not only in founder populations.

Although the present study relied on Quebec genealogies, the framework is not specific to genealogical data. It combines haplotype sharing, dimensionality reduction, and profile prediction, which can be applied in a wide range of populations. Deep genealogies provided an unusually detailed benchmark for predicting ancestry profiles, but other sources of information, including grandparental birthplace or geographic origin, could also be used to assess similar approaches. Future developments may further benefit from ancestral recombination graphs (ARGs), which offer a particularly rich representation of ancestry relationships.^48–51^

Several limitations should be noted. The genealogical completeness goes down with depth, so no band extends past the eighth generation, beyond which the record becomes too incomplete for training and validation.^52^ CARTaGENE recruited participants from the main urban centers of Quebec and so does not sample the province evenly, as volunteer cohorts generally do not.^53^ Every metric reported in this study is therefore computed on validation participants with no relative in the training set, so it should not be affected by the uneven sampling. Finally, the method depends on a register, and Balsac is essentially composed of Catholic parish records. A participant’s region of origin is therefore where their documented ancestors married, which is a historical and demographic designation rather than a biological one. Populations outside the register, such as Indigenous and non-Catholic communities, are underrepresented. Despite these limitations, the main conclusions remain consistent across the validation analyses.

Our results provide empirical support for recent calls to move beyond categorical population descriptors and toward richer representations of ancestry that better reflect the continuous nature of human genetic variation. Like any ancestry label, a single region of origin is a discrete category that may distort a continuous reality. A profile of proportions captures more documented ancestry than any single label could. Because the genome can predict these profiles where a register does not exist, it can serve as an indirect indication for individual demographic history. This demographic history is read not as ancestry from a single population but as a measured contribution from many ancestral sources. Collective efforts are required to move beyond labels and make more sensible descriptors available for all populations, including underrepresented ones.

## Data and code availability

Due to the informed consent given by study participants, CARTaGENE genotype data are available under restricted access via CARTaGENE’s independent Sample and Data Access Committee (SDAC, see *Web resources* below). Further information on access to CARTaGENE data is available in the article by McClelland et al.^19^ Access to the Balsac genealogical data is controlled in accordance with the Policy on Access to Balsac Data for Research Purposes to protect participant confidentiality and adhere to ethical guidelines. The procedure for requesting access to Balsac data is described on its data access page (see *Web resources* below). Access requests must be submitted to the Researchers Service of Projet Balsac and must include a completed Balsac Database Access Request Form along with all documents necessary for evaluation, such as a certificate of ethics approval. Applications are reviewed to ensure compliance with the scientific method, feasibility and Balsac access policies. Applicants are informed of the decision in writing. Questions regarding data access can be addressed to. If access is granted, data may be used only within the scope of the approved research project and must be destroyed once the authorized access period expires. The Balsac data are not publicly available due to ethical restrictions on the publication of genealogical information dating back less than 100 years.

The source code used for data processing, analysis and the production of figures in this study has been deposited in the following GitHub: https://github.com/Genopop/continuous_profiles.

## Acknowledgments

This work was made possible by the Digital Research Alliance of Canada which provided access to storage and computing resources. Funding for SLG was provided by the Canada Research Chair in Genetics and Genealogy CRC-2022-00444, of which he is the chair holder (https://www.chairs.gc.ca).

## Author contributions

All authors conceived and designed the study and drafted and approved the final version of the manuscript. GPM conceived the analysis pipeline and performed the analyses. CM and GPM played an important role in interpreting the results.

## Supplemental information

Supplemental information includes eight figures and two tables.

## Web resources

Balsac population register, https://balsac.uqac.ca

Balsac data access, https://balsac.uqac.ca/acces-donnees/

pbwt, https://github.com/richarddurbin/pbwt

GeneaKit, https://github.com/Genopop/geneakit

ibd_relatedness, S.R. Browning, https://faculty.washington.edu/sguy/ibd_relatedness.html

CARTaGENE, https://cartagene.qc.ca

CARTaGENE data access, https://cartagene.qc.ca/en/researchers/access-request.html

