## Supplementary material for "Continuous ancestry profiles from genomes and genealogies"

### Supplemental material

#### Comparison of the genome summaries

Before any model is fitted, genotypes need to be summarized into participant relatedness. We compared eight such summaries from the same genomes to determine how that choice affects the accurate prediction of regional ancestry.

Four summaries are derived from the observed-kinship (total IBD length) matrix: (1) the raw matrix itself, (2) a binary indicator of any nonzero entry, (3) the square root of each value, and (4) a logarithmic rescaling, $\ln(1 + \phi/\bar{\phi}^{+})$. Here $\bar{\phi}^{+}$ is the mean of non-zero off-diagonal entries among the participants with both genotype and genealogical data.

(5) A genotype relationship matrix (GRM) is commonplace in complex-trait analysis. Each variant is standardized by allele frequency, and the cross-product between participants is averaged across variants above 5% minor allele frequency. Its principal components routinely control for population structure,[1,2] following standard practice.

(6) A coancestry matrix from haplotype painting is the summary the analysis uses, obtained as described in the Material and methods.

(7) Spectral components of an IBD graph are a recently proposed alternative to the ordination (Material and methods). From our IBD segments, we built a graph joining any two participants whose total IBD sharing exceeds 6 cM, and used the leading components of that graph as coordinates in place of the principal coordinates.

(8) The combination of PBWTpaint and logarithmic IBD.

All eight followed the same protocol: the same regions and bands to predict, the same validation participants and the same cross-validation, using gradient boosting throughout, with one hundred axes each.

Haplotype painting scored better than the logarithm of observed kinship for the recent genealogical band, whereas the logarithm of observed kinship scored better for the intermediate and deep bands (Figure S1). Therefore, a combination of the logarithmic rescaling of observed kinship (Refined IBD) and of haplotype painting (PBWTpaint) was chosen for subsequent steps. Across the 1,850 validation participants, 50 principal coordinates of each ranked first, ahead of 100 principal coordinates of any separate summary. The differences remain significant after adjusting for multiple comparisons.

As a result, the GRM was the worst of the eight genetic summaries compared, whereas the haplotype- and IBD-based summaries were the best. In a young founder population such as Quebec, regional structure is better explained by recently co-inherited haplotypes better than by the allele frequencies of unlinked markers that a GRM measures.[3,4]

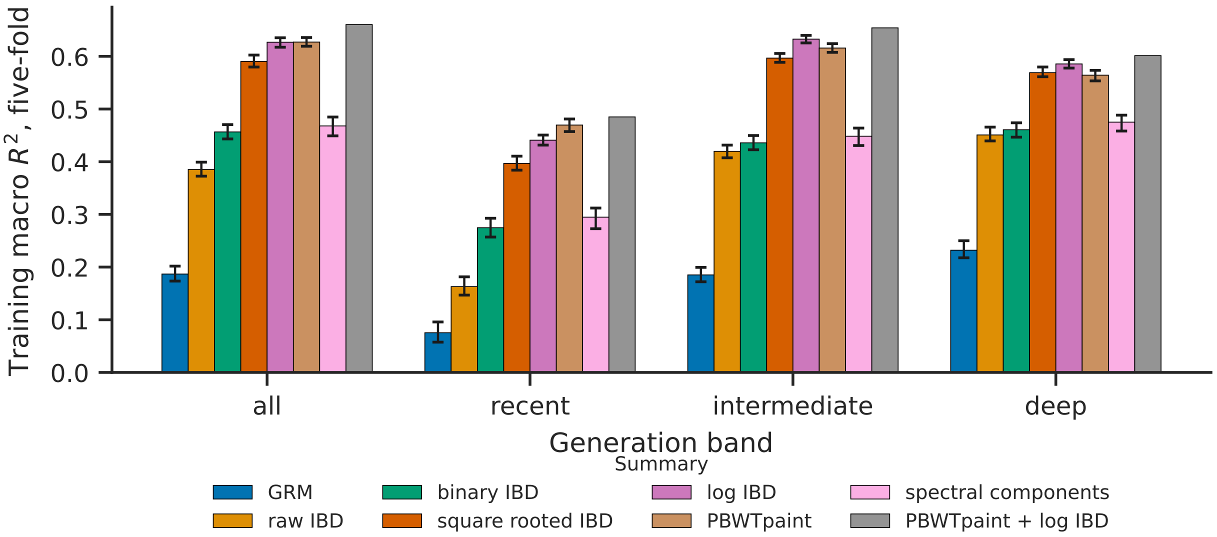


**Figure S1. How the genome is summarized affects ancestry recovery.** Gradient boosting was applied to eight inputs: the GRM, the raw IBD matrix, its binary, square-root and logarithmic rescalings, PBWTpaint coancestry, spectral components of the IBD graph, and a combination of PBWTpaint and logarithmic IBD. $R^2$ is the coefficient of determination between predicted and documented proportions for each region, averaged across the modeled regions. It is computed on all 1,850 validation participants. Error bars are 95% intervals of each input's difference from PBWTpaint and logarithmic IBD combined, the input the analysis uses, adjusted for 28 comparisons.

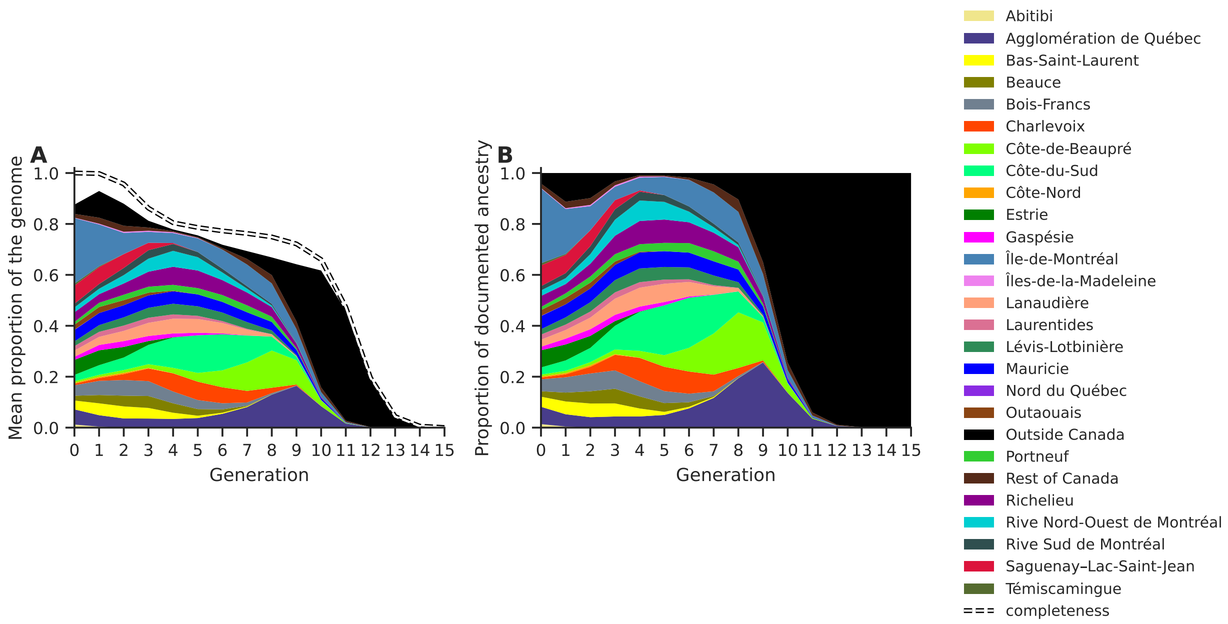


**Figure S2. Regional contributions change over time.** For each ancestor, the expected genetic contribution was computed and assigned to the region where their parents married. Panel (A) shows absolute contribution values. Totals decline as completeness goes down. Panel (B) presents the contributions as proportions of documented ancestry. At each generation, documented locations sum to one. Both estimates rely on ancestors of 9,405 CARTaGENE participants with both genotypes and genealogical information.

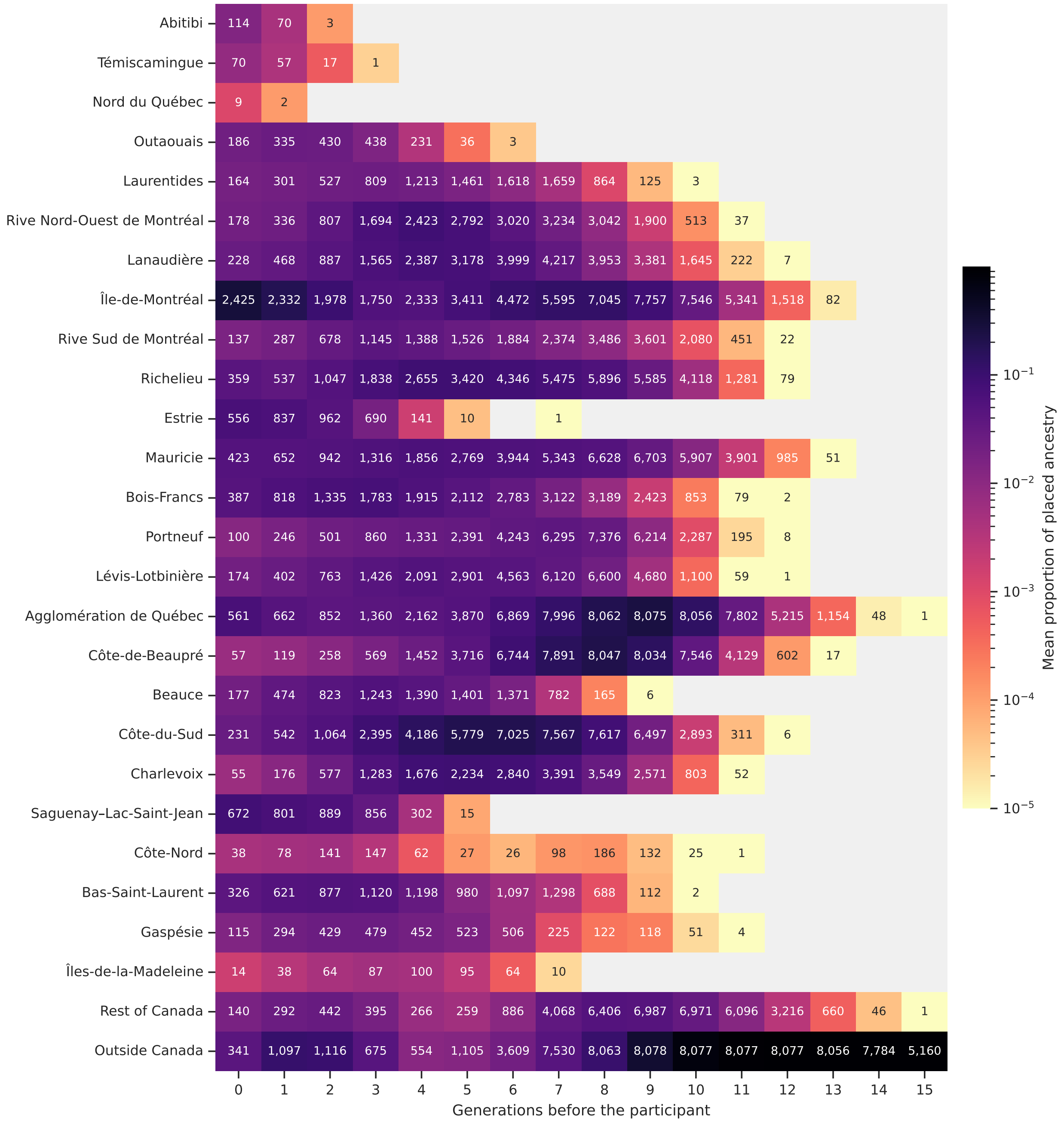


**Figure S3. Heat map of regional ancestry across generations.** Colors represent the mean ancestry proportion attributed to each region at each generation. Within a generation, proportions sum to one across regions. The number written in a cell is the count of participants holding ancestry in that region at that generation. A gray cell is a region holding none. An ancestor is attributed to the region where their parents married. Both are computed over the CARTaGENE participants with both genotype and genealogical data.

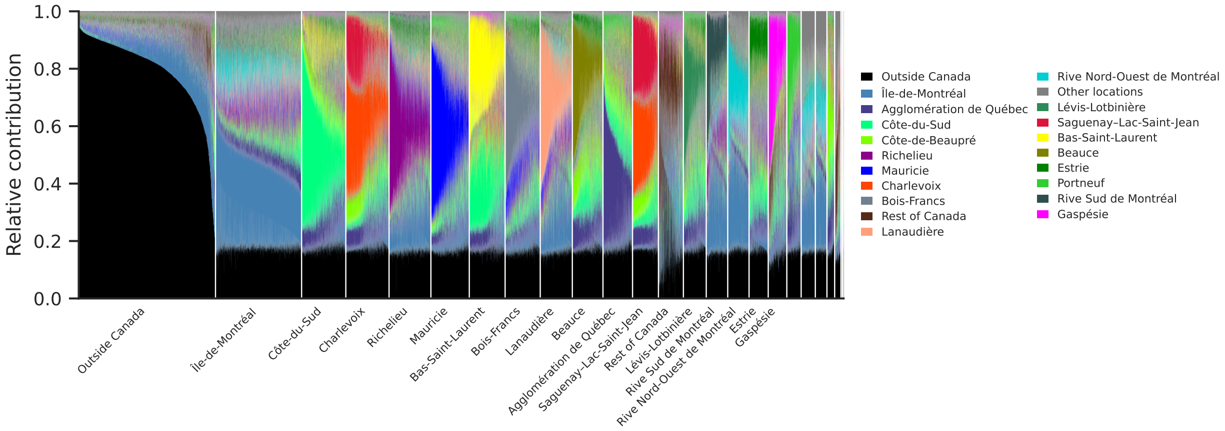


**Figure S4. Predicted ancestry profiles span several regions.** After regression was used to predict the ancestry proportion for each region from generation 0 to 15, the values were rescaled to sum to one. Each vertical bar represents one of the 28,332 CARTaGENE participants, and the vertical axis shows each region's predicted proportion.

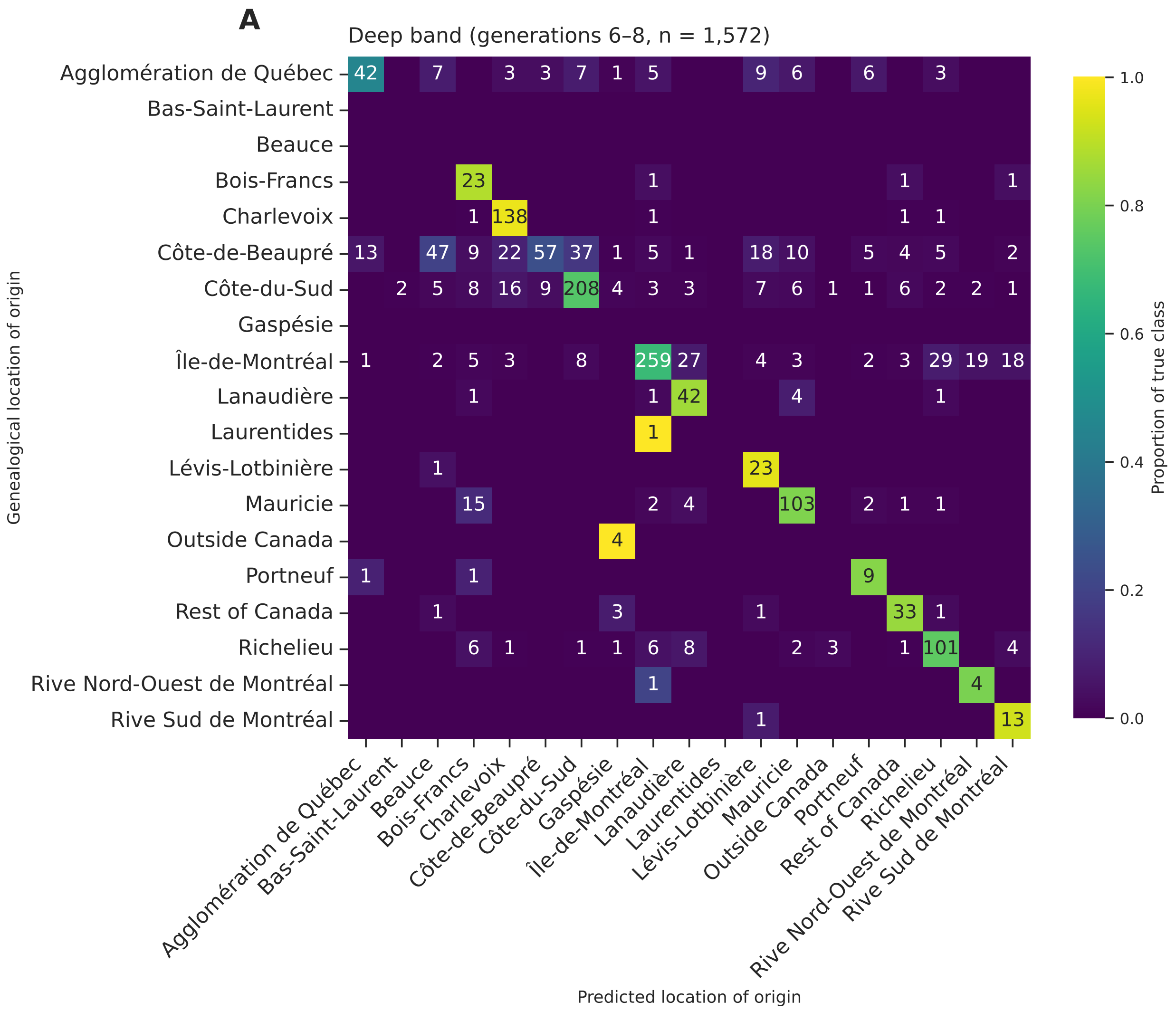
A
Deep band (generations 6–8, n = 1,572)
Genealogical location of origin
Predicted location of origin
Proportion of true class
Agglomération de Québec
Bas-Saint-Laurent
Beauce
Bois-Francs
Charlevoix
Côte-de-Beaupré
Côte-du-Sud
Gaspésie
Île-de-Montréal
Lanaudière
Laurentides
Lévis-Lotbinière
Mauricie
Outside Canada
Portneuf
Rest of Canada
Richelieu
Rive Nord-Ouest de Montréal
Rive Sud de Montréal

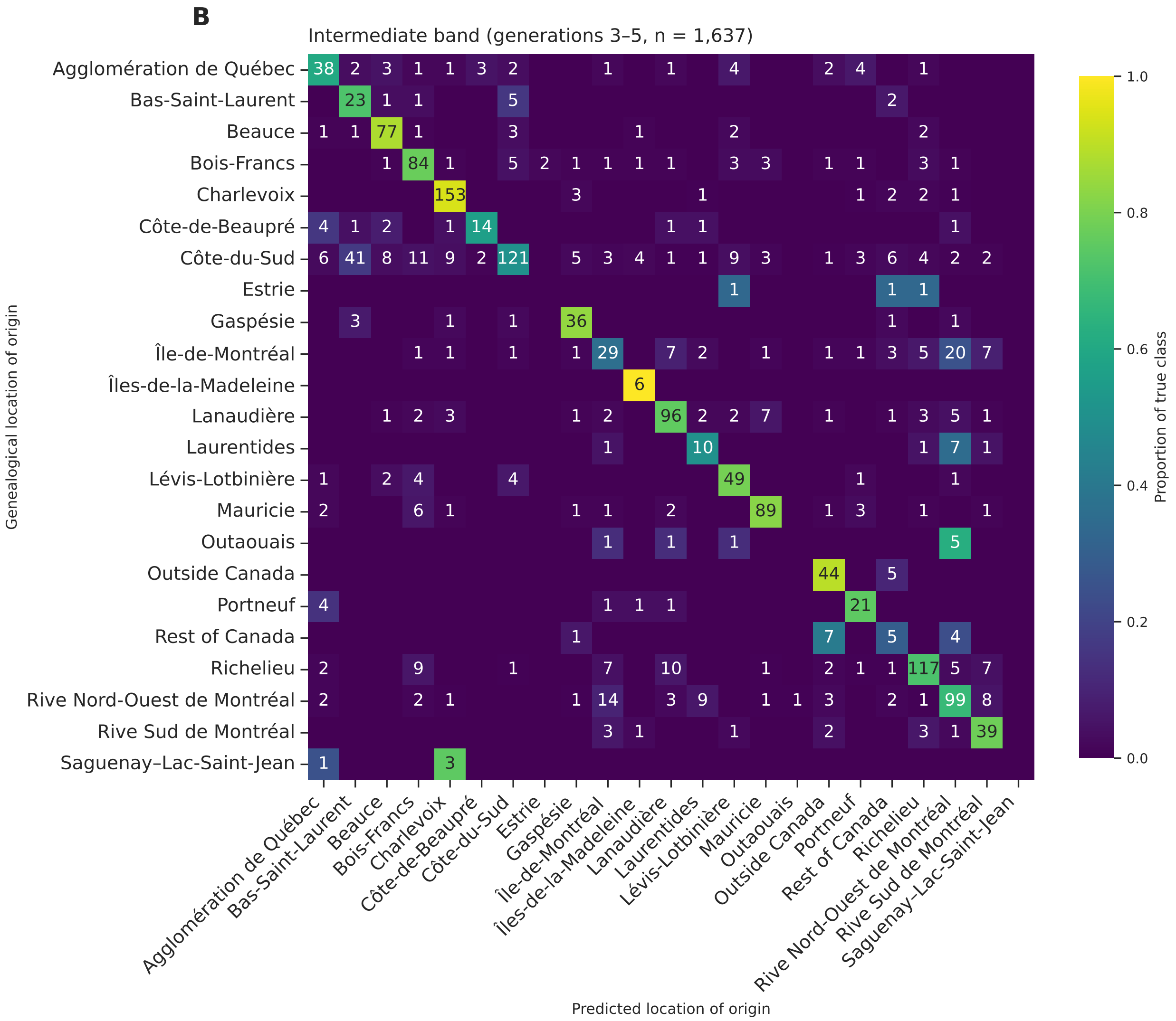
B
Intermediate band (generations 3–5, n = 1,637)
Genealogical location of origin
Predicted location of origin
Proportion of true class
Agglomération de Québec
Bas-Saint-Laurent
Beauce
Bois-Francs
Charlevoix
Côte-de-Beaupré
Côte-du-Sud
Estrie
Gaspésie
Île-de-Montréal
Îles-de-la-Madeleine
Lanaudière
Laurentides
Lévis-Lotbinière
Mauricie
Outaouais
Outside Canada
Portneuf
Rest of Canada
Richelieu
Rive Nord-Ouest de Montréal
Rive Sud de Montréal
Saguenay–Lac-Saint-Jean

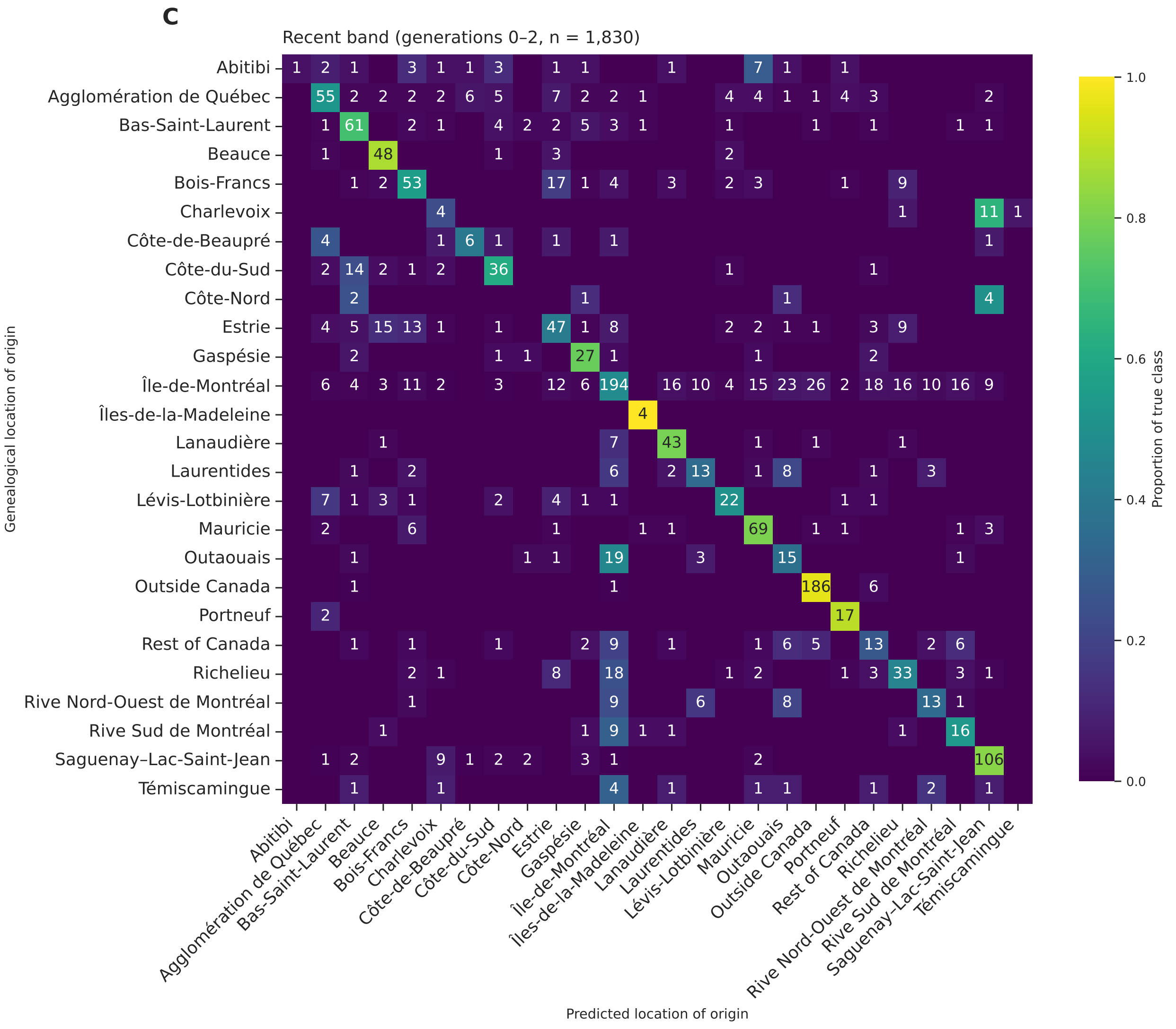


**Figure S5. Single-region prediction accuracy varies across time.** Confusion matrices showing the row-normalized accuracy of the classification for the validation participants with a documented region in each band: (A) generations 6–8 ($n$ = 1,572), (B) generations 3–5 ($n$ = 1,637) and (C) generations 0–2 ($n$ = 1,830).

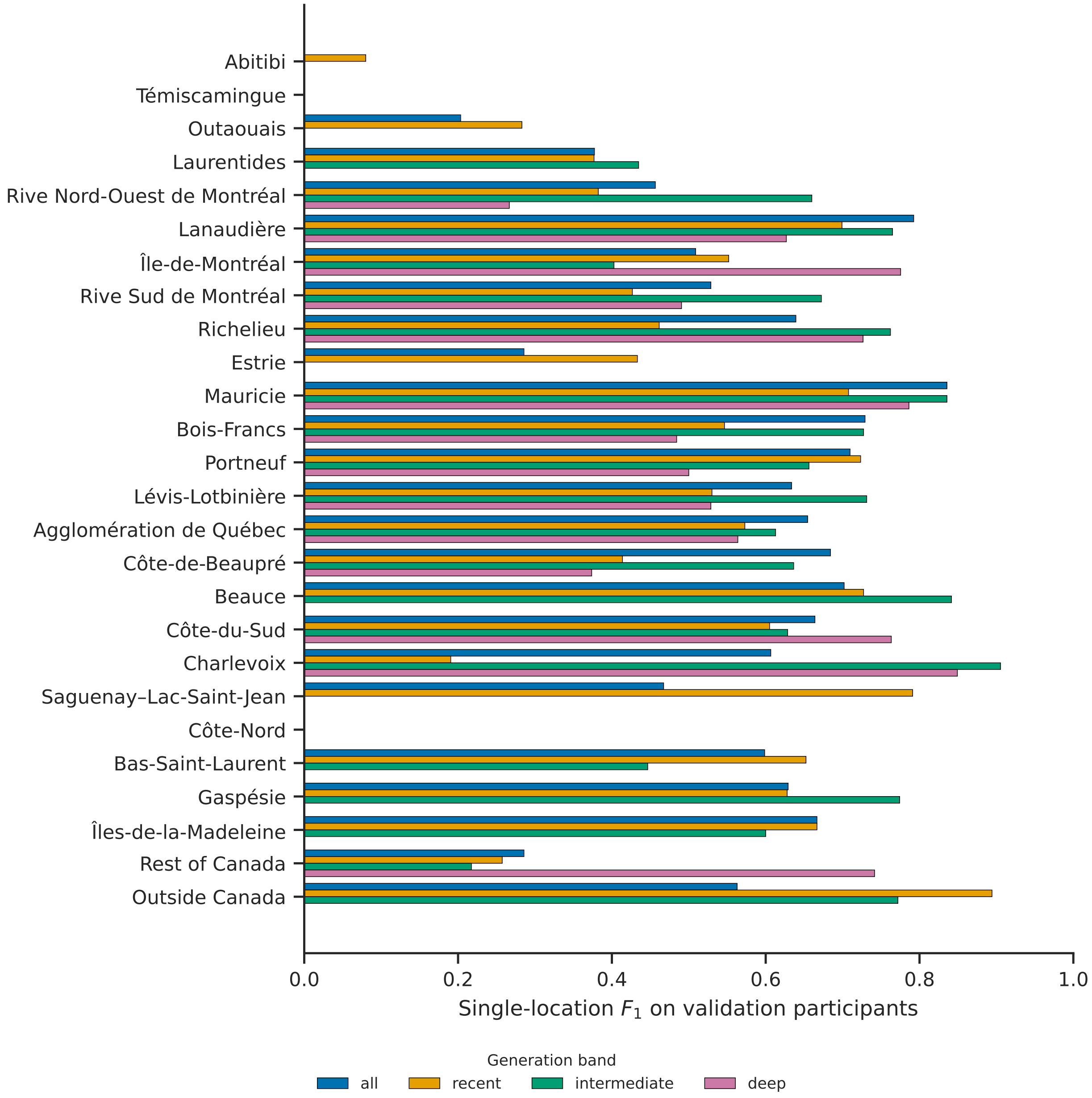


**Figure S6. The reliability of the predicted region of origin shifts across geography and generations.** Each participant was assigned the single region with the largest predicted proportion, after adjusting for how common that location is in the training set, as throughout the paper. An $F_1$ score, the harmonic mean of precision and recall, was then computed for each region against the largest documented region, one region at a time, among validation participants over all documented generations (generations 0–15, $n$ = 1,654) and in each band: recent (0–2, $n$ = 1,830), intermediate (3–5, $n$ = 1,637) and deep (6–8, $n$ = 1,572). The number of validation participants varies between bands because of incompleteness or undocumented marriage locations.

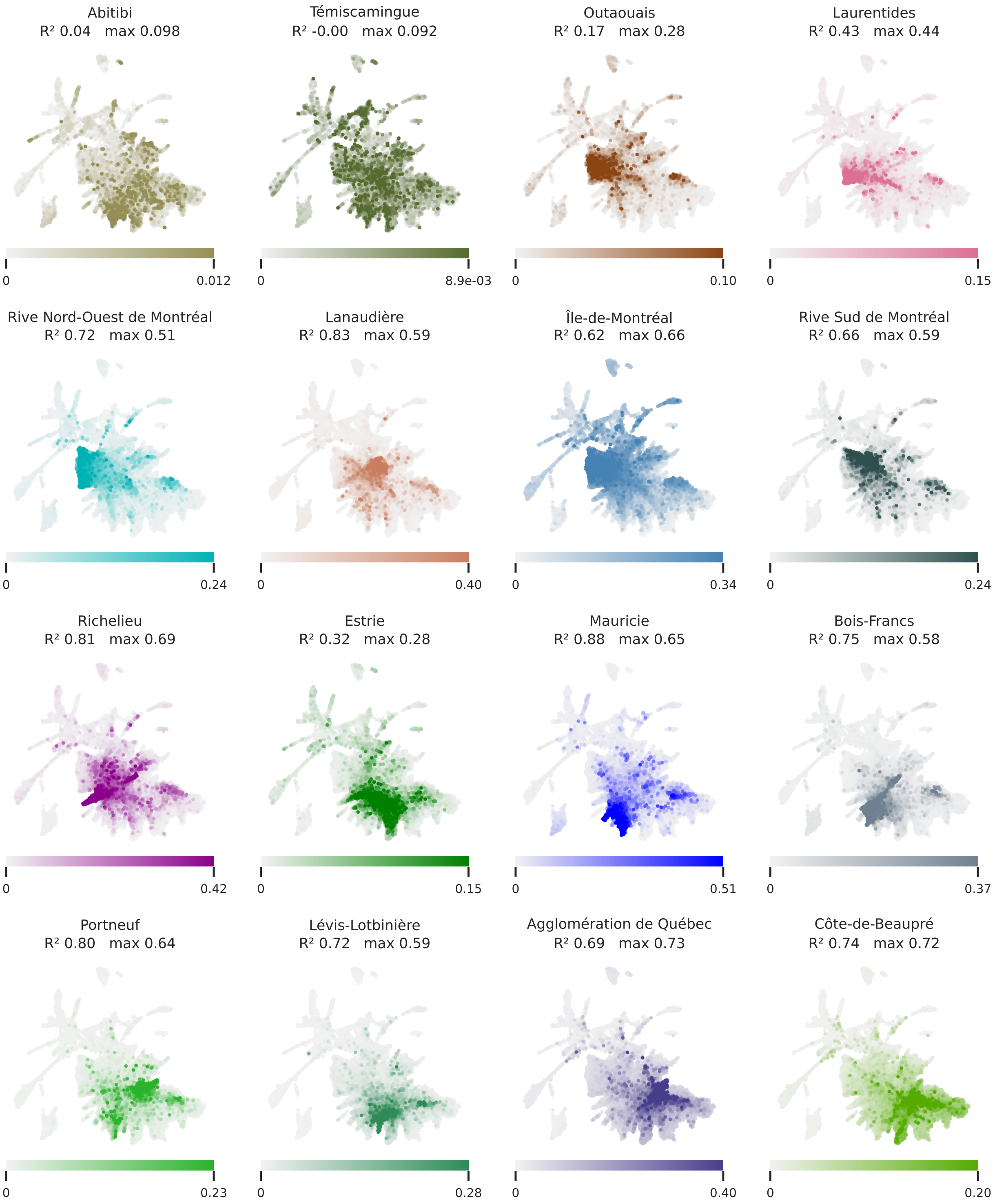

Abitibi
$R^2$ 0.04 max 0.098
0
0.012
Témiscamingue
$R^2$ -0.00 max 0.092
0
8.9e-03
Outaouais
$R^2$ 0.17 max 0.28
0
0.10
Laurentides
$R^2$ 0.43 max 0.44
0
0.15
Rive Nord-Ouest de Montréal
$R^2$ 0.72 max 0.51
0
0.24
Lanaudière
$R^2$ 0.83 max 0.59
0
0.40
Île-de-Montréal
$R^2$ 0.62 max 0.66
0
0.34
Rive Sud de Montréal
$R^2$ 0.66 max 0.59
0
0.24
Richelieu
$R^2$ 0.81 max 0.69
0
0.42
Estrie
$R^2$ 0.32 max 0.28
0
0.15
Mauricie
$R^2$ 0.88 max 0.65
0
0.51
Bois-Francs
$R^2$ 0.75 max 0.58
0
0.37
Portneuf
$R^2$ 0.80 max 0.64
0
0.23
Lévis-Lotbinière
$R^2$ 0.72 max 0.59
0
0.28
Agglomération de Québec
$R^2$ 0.69 max 0.73
0
0.40
Côte-de-Beaupré
$R^2$ 0.74 max 0.72
0
0.20

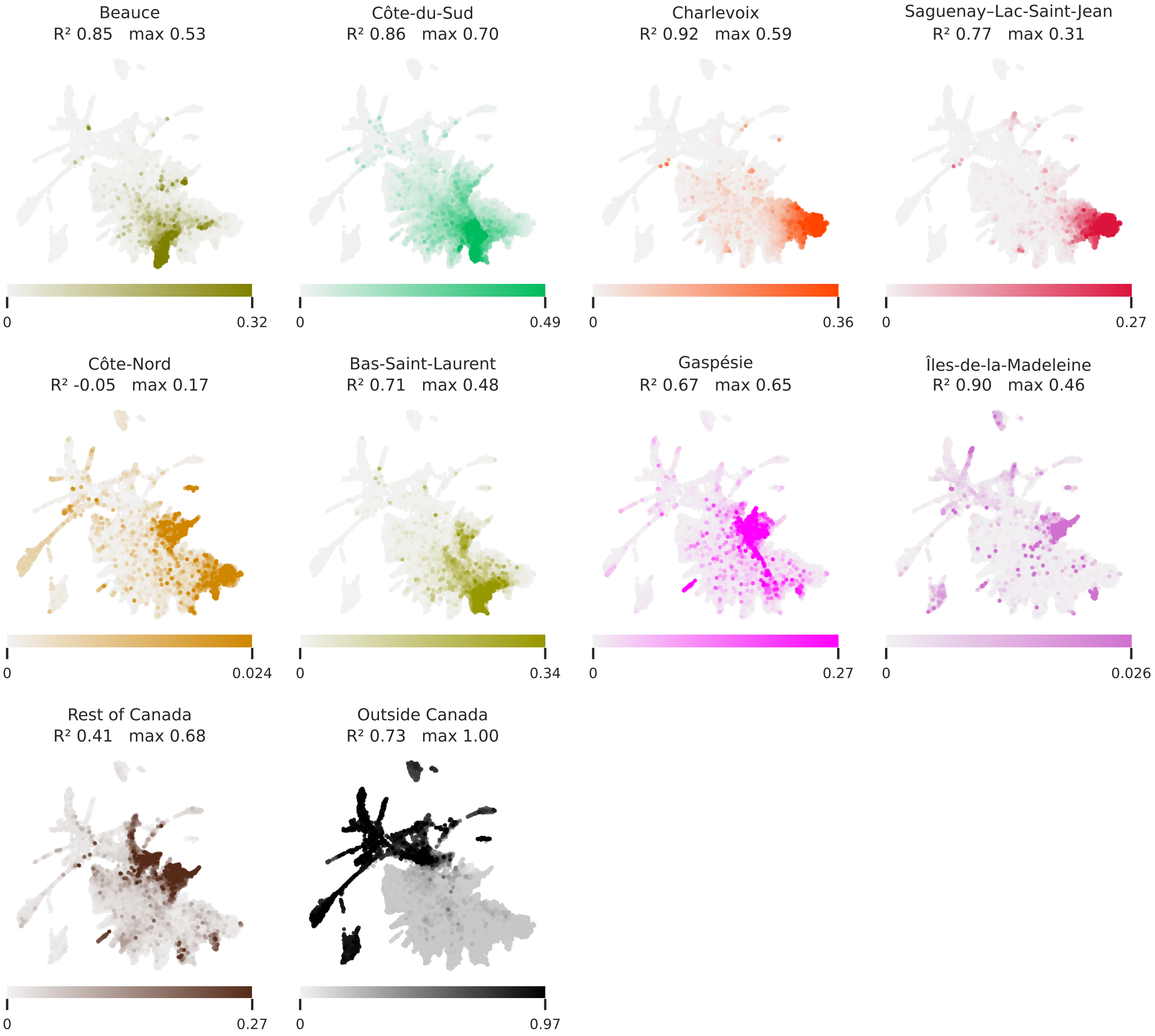


**Figure S7. Most participants show ancestry from multiple regions.** Each UMAP panel focuses on one region, using the same projection as Figure 4. Within a panel, color intensity reflects the region's predicted proportion for each person, with each panel scaled independently. $R^2$ is the coefficient of determination between predicted and documented proportions across 1,654 validation participants scored on all generations.

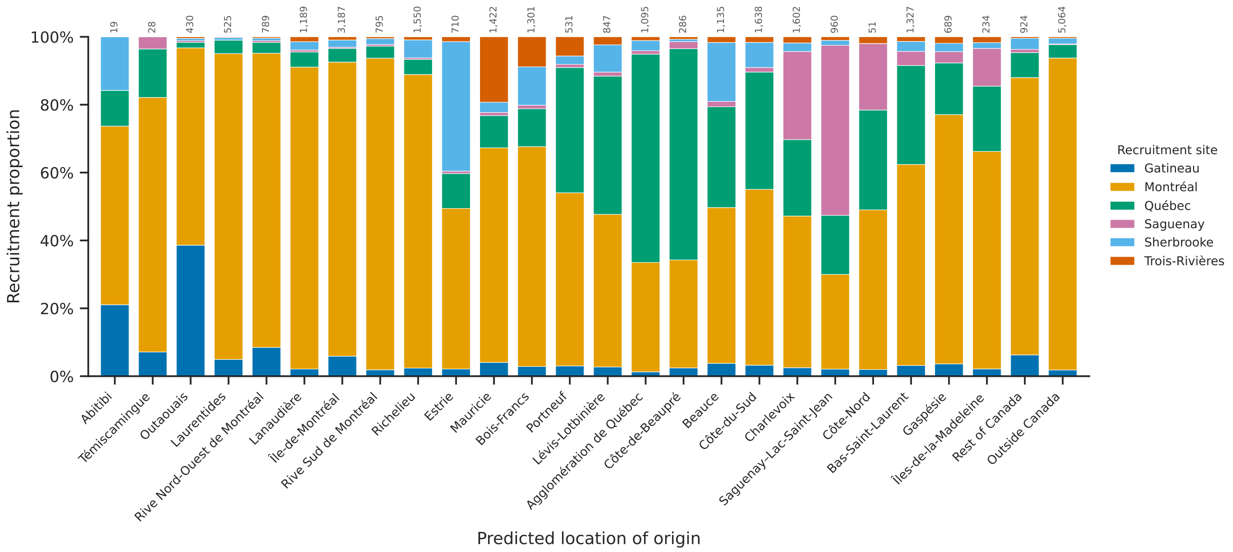


**Figure S8. Predicted regions are only weakly associated with the recruitment site.** For each predicted region of origin, the distribution of recruitment sites is shown for all 28,332 CARTaGENE participants assigned to that region by the classifier. Montreal is the most common recruitment site across nearly all predicted regions. Even so, regions with their own recruitment site, such as Saguenay–Lac-Saint-Jean and Mauricie, show a higher proportion of that site. Overall, recruitment location provides only a coarse approximation of regional ancestry.

**Table S1. Recovery by genealogical band.** Macro-$R^2$ is the coefficient of determination between predicted and documented proportions of ancestry for each region, averaged across regions common to the four genealogical bands. Ancestry placed in the correct region is the sum of the proportions of ancestry shared between predicted and documented ancestry for each region. The ancestry-blind baseline is the proportion of the profile predicted without any genetic relatedness information, or the average profile of the training set, that matches the documented profile. Those profile results are computed from one continuous value per region per participant. In contract, the single-region results are computed from one binary value per participant. The single-region accuracy is the proportion of participants whose predicted dominant region of origin corresponds to their actual documented region of origin. The majority-label location corresponds to the most common region in the training participants. The majority-label baseline corresponds to the single-region accuracy if that location was always predicted by the model. Due to the different nature of whole profiles and single-region accuracy, they can not be directly compared.

| | **Recent (0–2)** | **Intermediate (3–5)** | **Deep (6–8)** | **All (0–15)** |
|---|---|---|---|---|
| **Whole profile** | | | | |
| Macro-$R^2$ | 0.50 | 0.67 | 0.58 | 0.68 |
| Ancestry placed in correct region | 57% | 72% | 83% | 78% |
| Ancestry-blind baseline | 15% | 27% | 55% | 48% |
| **Single-region** | | | | |
| Single-region accuracy | 59% | 70% | 67% | 60% |
| Majority-label baseline | 22% | 15% | 24% | 20% |
| Majority-label location | Île-de-Montréal | Côte-du-Sud | Île-de-Montréal | Île-de-Montréal |
| Locations as largest documented region | 26 | 23 | 16 | 26 |

**Table S2. Regional distribution of the CARTaGENE cohort for each recruitment site.** The recruitment site is found in columns and the region where a participant's parents married (generation 0 of the genealogy) is found in rows. Four individuals lack a recorded recruitment site and are omitted.

| | Recruitment site | | | | | | |
|---|---|---|---|---|---|---|---|
| **Parents' marriage region** | **Gatineau** | **Montréal** | **Québec** | **Saguenay** | **Sherbrooke** | **Trois-Rivières** | **Total** |
| Abitibi | 17 | 73 | 9 | 4 | 10 | 1 | 114 |
| Témiscamingue | 8 | 49 | 6 | 3 | 2 | 2 | 70 |
| Nord du Québec | 2 | 5 | 1 | 1 | 0 | 0 | 9 |
| Outaouais | 100 | 77 | 5 | 1 | 2 | 1 | 186 |
| Laurentides | 8 | 147 | 7 | 0 | 1 | 1 | 164 |
| Rive Nord-Ouest de Montréal | 5 | 167 | 4 | 0 | 1 | 1 | 178 |
| Lanaudière | 5 | 195 | 13 | 0 | 12 | 3 | 228 |
| Île-de-Montréal | 47 | 2,220 | 80 | 6 | 50 | 22 | 2,425 |
| Rive Sud de Montréal | 3 | 120 | 6 | 0 | 7 | 1 | 137 |
| Richelieu | 11 | 317 | 22 | 1 | 5 | 3 | 359 |
| Estrie | 15 | 222 | 52 | 3 | 262 | 2 | 556 |
| Mauricie | 16 | 226 | 52 | 2 | 12 | 115 | 423 |
| Bois-Francs | 16 | 256 | 53 | 1 | 32 | 29 | 387 |
| Portneuf | 1 | 39 | 52 | 1 | 4 | 3 | 100 |
| Lévis-Lotbinière | 1 | 67 | 96 | 3 | 5 | 2 | 174 |
| Agglomération de Québec | 15 | 159 | 373 | 3 | 7 | 4 | 561 |
| Côte-de-Beaupré | 1 | 19 | 35 | 2 | 0 | 0 | 57 |
| Beauce | 6 | 75 | 72 | 3 | 18 | 3 | 177 |
| Côte-du-Sud | 3 | 95 | 110 | 5 | 14 | 4 | 231 |
| Charlevoix | 2 | 25 | 22 | 5 | 1 | 0 | 55 |
| Saguenay—Lac-Saint-Jean | 19 | 192 | 106 | 341 | 10 | 4 | 672 |
| Côte-Nord | 1 | 17 | 16 | 4 | 0 | 0 | 38 |
| Bas-Saint-Laurent | 19 | 189 | 100 | 8 | 5 | 5 | 326 |
| Gaspésie | 8 | 81 | 20 | 2 | 4 | 0 | 115 |
| Îles-de-la-Madeleine | 0 | 6 | 5 | 1 | 2 | 0 | 14 |
| Rest of Canada | 28 | 88 | 15 | 1 | 6 | 2 | 140 |
| Outside Canada | 8 | 294 | 29 | 1 | 7 | 2 | 341 |
| Other or unknown | 715 | 14,313 | 2,969 | 772 | 853 | 469 | 20,091 |
| **Total** | **1,080** | **19,733** | **4,330** | **1,174** | **1,332** | **679** | **28,328** |
